# Integrated Analysis of T cell Repertoire and Transcriptome Identifies Mechanisms of Regulatory T cell (Treg) Suppression of Acute Graft-versus-Host-Disease

**DOI:** 10.1101/2022.07.26.501553

**Authors:** Juliane K. Lohmeyer, Toshihito Hirai, Mustafa Turkoz, Stephane Buhler, Teresa Lopes Ramos, Natalie Köhler, Jeanette Baker, Xuhuai Ji, Jean Villard, Yves Chalandon, Federico Simonetta, Robert S. Negrin

## Abstract

CD4+FOXP3+ regulatory T cells have demonstrated efficacy in graft-versus-host disease (GvHD) prevention and treatment. Preclinical and clinical studies indicate that Treg are able to protect from GvHD without interfering with the graft-versus-tumor (GvT) effect of hematopoietic cell transplantation (HCT), although the underlying molecular mechanisms are largely unknown. To elucidate Treg suppressive function during *in vivo* suppression of acute GvHD, we performed paired T cell receptor (TCRα, TCRβ genes) repertoire sequencing and RNA sequencing analysis on conventional T cells (Tcon) and Treg before and after transplantation in an MHC major-mismatch mouse model of HCT. We show that both Treg and Tcon underwent clonal restriction and that Treg did not interfere with the activation of alloreactive Tcon clones and the breadth of their TCR repertoire, however, markedly suppressed their expansion. Transcriptomic analysis revealed that Treg predominantly affected the transcriptome of CD4 Tcon and to a lesser extent of CD8 Tcon, modulating the transcription of genes encoding pro- and anti-inflammatory molecules as well as enzymes involved in metabolic processes, inducing a switch from glycolysis to oxidative phosphorylation. Finally, Treg did not interfere with the induction of gene sets involved in the GvT effect. Our results shed light into the mechanisms of acute GvHD suppression by Treg and will support the clinical translation of this immunoregulatory approach.

**Key Points:**

- Regulatory T cells modulate conventional T cells transcriptome during GvHD suppression by affecting several, non-redundant pathways.
- Regulatory T cells undergo activation and clonal expansion during GvHD suppression.

## Introduction

Allogeneic hematopoietic cell transplantation (HCT) is a well-established and potentially curative therapy for a broad range of hematologic malignancies due to the graft-versus-tumor (GvT) effect. Unfortunately, allogeneic HCT is still associated with significant morbidity and mortality related to cancer relapse and transplant complications, namely graft-versus-host Disease (GvHD). The immunological mechanism responsible for GvHD, i.e. donor T cell alloreactivity toward host antigens, is also responsible for the beneficial GvT effect of allogeneic HCT^1^. Because of the interconnection between these two phenomena, none of the currently employed strategies to GvHD prevention and treatment can efficiently target GvHD without affecting GvT.

Immunoregulatory cellular therapies are a promising approach for GvHD prevention and treatment^2,3^. We and others have previously shown in preclinical murine models that CD4+FOXP3+ regulatory T cells (Treg) are able to protect from GvHD without interfering with the GvT effect of HCT^4–6^. Efforts are ongoing to translate the use of Treg adoptive transfer for GvHD prevention^7–10^ and treatment^11,12^ to the clinic with promising results.

The precise cellular and molecular mechanisms underlying GvHD suppression by Treg are incompletely understood. Two non-exclusive and potentially complementary models exist: Treg could quantitatively affect T cell responses by limiting the activation and expansion of alloreactive T cell clones and/or might qualitatively modulate T cell function by selectively interfering with pathways responsible for GvHD but dispensable for GvT. To gain further insights supporting one or the other of these models, we performed paired T cell receptor (TCR) and RNA sequencing analysis on Tcon and Treg before and after transplantation using an MHC major-mismatch mouse model of acute GvHD.

## Material and methods

### Animals

Eight-to 12-week-old BALB/cJ (H-2kd), CD45.2 Thy1.2 C57Bl/6J (H-2kb) mice and CD45.1 Thy1.2 C57Bl/6 (B6.SJL-*Ptprc^a^ Pepc^b^*/BoyJ) mice were purchased from Jackson Laboratory. C57Bl/6 Foxp3GFP-DTR mice were provided by Alexander Rudensky (Memorial Sloan Kettering Cancer Center, New York, NY). Firefly luciferase (Luc)+ transgenic CD45.1 Thy1.1 C57Bl/6 L2G85 mice have been described previously^13^. All experiments performed on animals were approved by Stanford University’s Institutional Animal Care and Use Committee and were in compliance with the guidelines of humane care of laboratory animals.

### Acute GvHD murine model

Donor CD4^+^ and CD8^+^ conventional T cells (Tcon) were separately isolated from splenocytes harvested from CD45.1 Thy1.1 luc^+^ C57Bl/6 mice by negative enrichment (Stemcell). T cell–depleted bone marrow (TCD-BM) cells were prepared from CD45.1 Thy1.2 C57Bl/6 mice by first crushing bones followed by T cell depletion using CD4 and CD8 MicroBeads (Miltenyi Biotec). CD45.2 Thy1.2 FoxP3/GFP^+^ CD4^+^ Treg were FACS sorted from CD8/CD19-depleted single cell suspensions from spleens and lymph nodes from CD45.2 Thy1.2 FoxP3GFP^+^ C57Bl/6 using a BD FACS Aria II. CD45.2 Thy1.2 BALB/c mice were lethally irradiated (8.8 Gy) and transplanted with 5×10e6 TCD-BM cells from CD45.1 Thy1.2 C57Bl/6 mice alone or together with CD45.2 Thy1.2 C57Bl/6 FoxP3/GFP^+^ Treg (1×10e6) on day 0. On day 2, CD45.1^+^ Thy1.1^+^ C57Bl/6 Tcon (1×10e6; CD4:CD8 ratio = 2:1) were injected to induce GvHD. Irradiated (11 Gy) syngeneic C57Bl/6 recipients receiving C57Bl/6 CD45.1^+^ Thy1.2^+^ TCD-BM and CD45.1^+^ Thy1.1^+^ Tcon alone were used as controls. Mice were monitored daily, and body weight and GvHD score were assessed weekly.

### BLI

Bioluminescent imaging (BLI) was performed as previously described^14^ on day 8 after HCT. Briefly, mice were injected with D-Luciferin Firefly (Biosynth; 10 mg/kg, i.p.) intraperitoneally 10 minutes before being anesthetized with 2% isoflurane in oxygen. Acquisition was performed using an Ami Imager, and images were analyzed with Aura software (Spectral Instruments Imaging, Tucson, AZ).

### Cell isolation

Recipient mice were euthanized 8 days after HCT (6 days after Tcon adoptive transfer) and single cell suspensions obtained from spleens and lymph nodes. Cells from 3 animals per group were pooled to obtain 2 biological replicates in each of the two independent experiments. After Fc block (Miltenyi), cells were incubated with the following antibodies (BioLegend): CD4 (BV421), CD8 (BV605), Thy1.1 (PE), CD45.1 (PE-Cy7), CD45.2 (APC), H-2kd (biotin) and CD19 (biotin) followed by streptavidin APC-Fire. Donor-derived Thy1.1^+^ CD45.1^+^ CD4 and CD8 Tcon as well as Thy1.2^+^ CD45.2^+^ CD4^+^ FoxP3/GFP^+^ Treg were FACS-sorted. T cell subsets before injection and recovered at day 8 post-HCT were frozen in Trizol (Thermo Fisher Scientific) and conserved at −80°C until analysis.

### RNA sequencing analysis

RNA was extracted using the TRizol RNA isolation method (Cat. #15596026, Thermo-fisher Scientific, USA) combined with the RNeasy MinElute Cleanup (Cat. #74204, QIAGEN, Germany). SMARTseq v4 Ultra Low Input RNA kit for Sequencing (Cat. # 634890, Takara Bio, USA) was used to generate full-length cDNA prior to library preparation with the Nextera XT DNA Library Prep Kit (Illumina, Inc.). After pooling, libraries were sequenced on the Illumina HiSeq 4000 platform (75 bp, paired-end). Sequencing reads were checked using FastQC v.0.11.7. The pseudo-aligner Kallisto^15^ was used to estimate transcript counts and transcripts per million (TPM) for the mouse genome assembly GRCm38 (mm10). Transcript-level abundance was quantified and summarized into gene level using the tximport R package. Differential gene expression was performed using the DESeq2 R package version 1.22.221, using FDR < 0.05. Heatmaps were generated using Pheatmap version 1.0.12., Principal component analysis (PCA) was performed using the FactoMineR package version 1.41 and visualized using the factorextra package version 1.0.5. For volcano plots, packages EnhancedVolcano version 1.14.0 and apeglm version 1.18.0 were used. Gene-set enrichment analysis was conducted using the fgsea R package.

### TCR sequencing analysis

For TCR sequencing, libraries were prepared from the synthesized full-length cDNA using the nested PCR method reported previously^16,17^, with a modification to amplify bulk TCRα and TCRβ repertoire genes.. Sequencing was performed using the Illumina MiSeq platform after Illumina paired-end adapters incorporation. TCRα and TCRβ sequence analysis was performed with VDJFasta. After total count normalization, downstream analysis was performed as previously described^18^. Briefly, the Shannon clonality index was used to estimate T cell repertoire diversity, with zero defining maximum diversity (i.e., polyclonal samples) and one minimum diversity (i.e., monoclonal samples). TCIRα and TCRβ repertoire overlap analyses were based on the amino acid sequences of the CDR3 region and calculated using the Jaccard index (number of shared clonotypes between two samples divided by the total number of clonotypes in both samples). This index measures the overlap based on the statistical dispersion of clonotypes in the samples and is expected to vary between 0 (no similarity) and 1 (complete similarity). Scatterplots, barplots, and boxplots were generated using R version 3.5.1.

### Statistical analysis

The Mann–Whitney U test was used in cross-sectional analyses to determine statistical significance. Survival curves were represented with the Kaplan–Meier method and compared by log-rank test. Statistical analyses were performed using R version 3.5.1 Comprehensive R Archive Network (CRAN) project (http://cran.us.r-project.org) with R studio version 1.1.453.

## Results

### Treg treatment inhibited Tcon expansion and target tissue infiltration without affecting their TCR repertoire breadth

We employed a well-established mouse model of GvHD in which Treg were transferred at time of HCT, two days prior the adoptive transfer of Tcon^19^. As previously described, mice treated with Treg prior to Tcon transfer showed significantly improved survival and GvHD scores compared to mice receiving Tcon alone (Supplemental Figure 1A). The use of Tcon isolated from luciferase^+^ donors revealed that this effect was associated with a significant reduction of Tcon expansion at day 8 after transplantation (day 6 after Tcon administration; Supplemental Figure 1B-C). At this time point, previously reported to be the peak of Tcon expansion^13^, Treg treatment also affected the localization of luc^+^ Tcon. The Tcon derived signal was mainly restricted to secondary lymphoid organs (spleen and lymph nodes) and reduced in the abdominal region in mice receiving Treg compared to untreated mice with GVHD (Supplemental Figure 1B).

Based on these results, we first hypothesized that Treg would inhibit the expansion of alloreactive clones during GvHD by controlling the TCR repertoire breadth, similarly to what was previously shown during antiviral responses^20^. To this aim, at day 8 after HCT, we re-isolated donor-derived CD45.1^+^ Thy 1.1^+^ CD4 and CD8 Tcon previously administered to syngeneic CD45.2^+^ C57Bl/6 mice or allogeneic CD45.2^+^ BALB/c mice in the presence or absence of CD45.2^+^ FOXP3^gfp+^ Treg (Figure 1A). As expected, the analysis of the TCR repertoire based on sequencing of the TCR alpha and beta chains revealed a significant clonal restriction of both CD4 (Figure 1B, left panel) and CD8 (Figure 1C, left panel) Tcon recovered from allogeneic HCT recipients compared with Tcon from syngeneic HCT recipients. Importantly, such clonal restriction in allogeneic HCT recipients was not inhibited by Treg treatment (Figure 1B-C, left panels). Clonal overlap between Tcon collected at day 8 and those before injection was reduced in allogeneic HCT recipients compared to syngeneic controls (Figure 1B-C, middle panels) and Treg did not inhibit such reduction in clonal overlap (Figure 1B-C, right panels). Collectively, our data indicate that Treg affected the expansion and the localization of Tcon after HCT without impacting the TCR repertoire breadth of Tcon and the initial activation of alloreactive T cell clones during GvHD.

**Figure 1.**
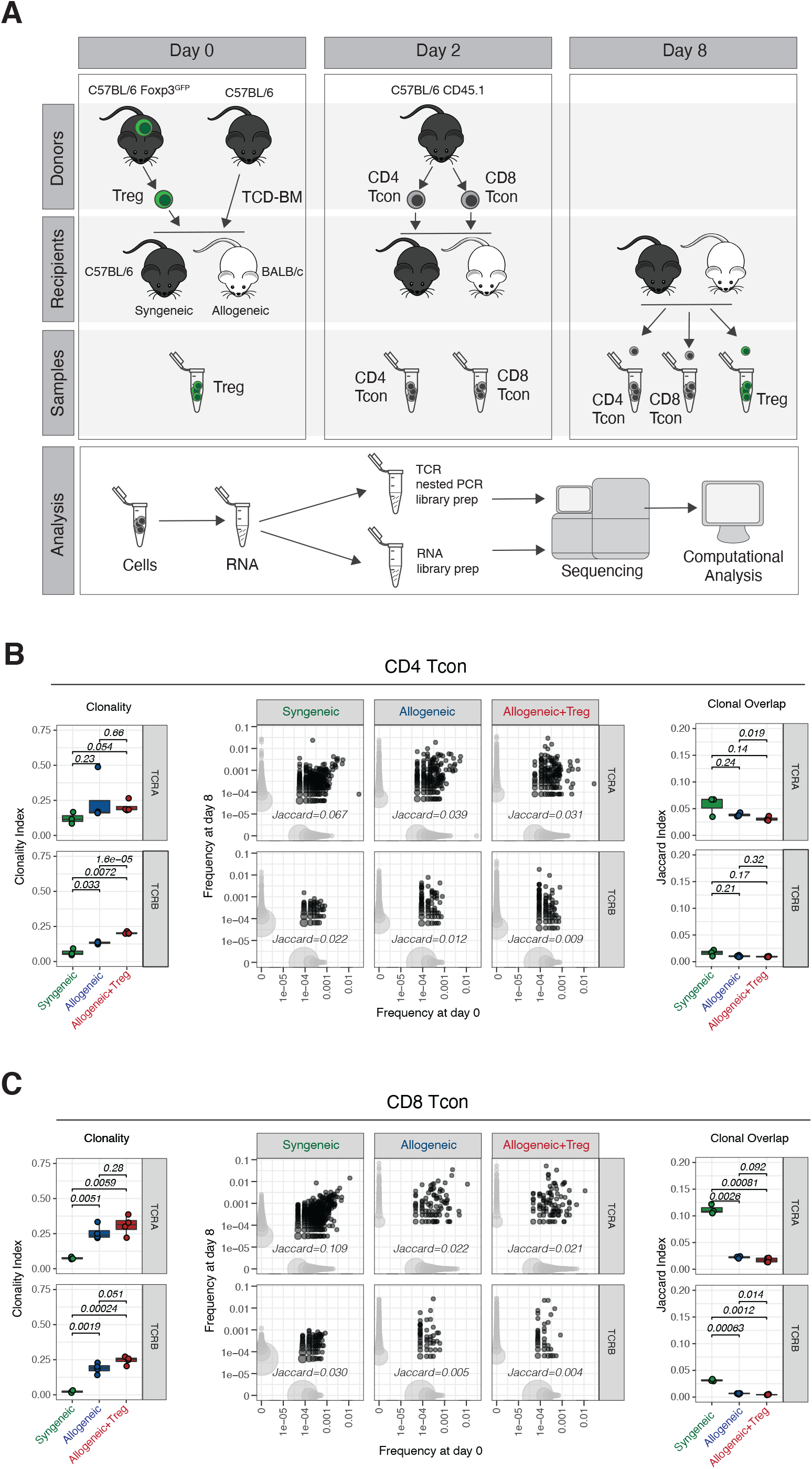
Treg did not affect the TCR breadth of Tcon after HCT. (A) Schematic representation of the experimental pipeline. On day 0, Balb/c or C57Bl/6 recipient mice were lethally irradiated and transplanted with 5×10^6^ CD45.1^+^ Thy 1.2^+^ T cell depleted bone marrow cells (TCD-BM) with or without 1×10^6^ Foxp3^GFP^ Treg cells from C57Bl6 donors. On day 2, 1×10^6^ CD45.1^+^Thy1.1^+^ Tcon from C57Bl/6 donors were injected. GFP^+^ donor Treg and CD45.1^+^ Thy1.1^+^ CD4 and CD8 donor Tcon before transplantation (day 0/2) and isolated on day 8 were used for sequencing analysis (B, C) Clonality of the TCRA and TCRB repertoire in CD4 (B) and CD8 (C) Tcon recovered at day 8 after HCT in syngeneic recipients (green box and symbols), allogeneic recipients (blue box and symbols) and allogeneic recipients receiving Treg (red box and symbols). Representative example of overlap of the TCRA and TCRB repertoire in CD4 and CD8 Tcon prior to transplantation and at day 8 after HCT (left panels). Scatter plots (middle panels) represent clone frequencies before and after HCT and number of unique clones (dot size). Clones that are only observed at one time point are colored in light grey, while overlapping clones are colored in dark grey. Repertoire overlap in CD4 and CD8 (right panels) Tcon recovered at day 8 after HCT and before injection quantified using the Jaccard index of similarity. Groups were compared using a nonparametric Mann–Whitney U test and p values are shown.

### Treg treatment affected CD4 and to a lesser extent CD8 Tcon transcriptome during GvHD

We next evaluated the impact of Treg treatment on CD4 and CD8 Tcon at the transcriptomic level. Principal component analysis (PCA) of the top 1000 most differentially expressed genes across all samples revealed that 68% of the variance was explained by PC1, which clearly segregated CD4 and CD8 Tcon recovered at day 8 from allogeneic recipients from cells before injection or recovered from syngeneic recipients (Figure 2A), revealing a dominant effect of the allogeneic transplant procedure on the different T cell populations. PC1 was mainly driven by naïve T cell genes (*Ccr7*, *Sell*, *Il6ra*, *Il6st*, *Foxo1*) that were progressively downregulated along PC1, pointing to T cell activation/effector differentiation as a main element affected by the transplantation into allogeneic mice and, to a lesser extent, into syngeneic recipients (Figure 2A). Treg impact on CD4 and CD8 Tcon transcriptome was revealed by PC2 (Figure 2A) which contributed to 14.8% and 17.3% of the variance in CD4 and CD8 Tcon, respectively. Treg treatment mainly affected CD4 Tcon transcriptome (Figure 2A, left panel) inducing the downregulation of 219 genes and the upregulation of 111 genes (Figure 2B, left panel) compared to Tcon in the absence of Treg. In particular, Treg induced the downregulation of Th1-signature genes (*Tbx21*, *Il12rb1*, *Il12rb2*) and proinflammatory genes (*Il18rap*) while promoting up-regulation of anti-inflammatory genes (*Il18bp*) and Th2 signature genes (*Ccr4*, *Il4*) in CD4 Tcon (Figure 2B left panel, Supplemental Figure 2A). Conversely, only a limited impact of Treg treatment was observed on CD8 Tcon (Figure 2A, right panel) with only 19 genes upregulated and 17 genes downregulated (Figure 2B, right panel) in Treg-treated CD8 Tcon compared to untreated CD8 Tcon. Collectively, these results revealed that Treg did not interfere with the major transcriptomic changes associated with T cell activation/effector differentiation during GvHD but exerted a CD4-dominant immunomodulatory effect on lineage-specific genes suppressing Th1 differentiation.

**Figure 2.**
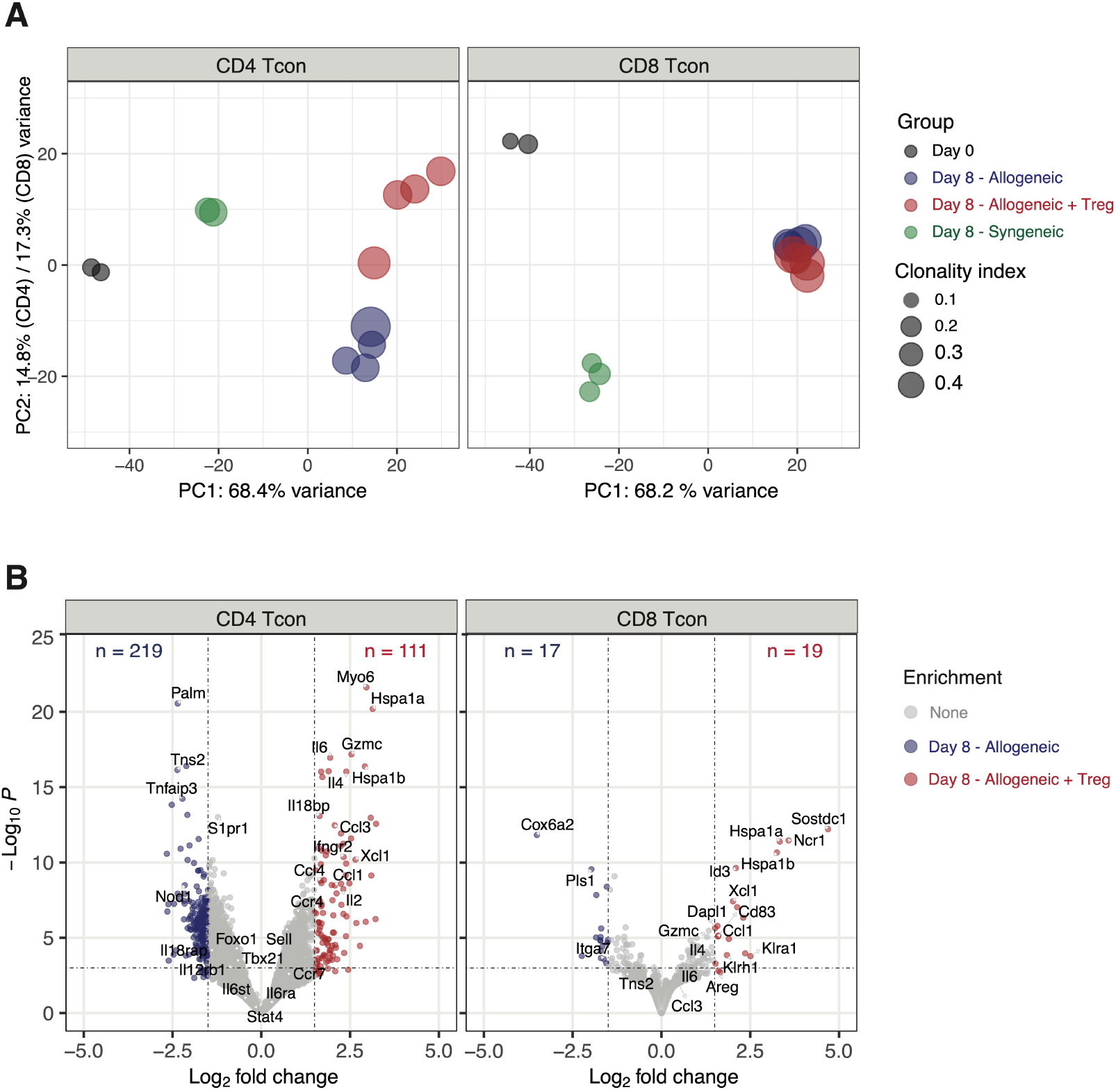
Treg modulated CD4 and to a lesser extent CD8 Tcon transcriptome during GvHD. (A) Principal component analysis of transcriptome based on the top 1000 differentially expressed genes across all CD4 (left panel) and CD8 (right panel) Tcon samples. (B) Volcano plots showing significance and Log2 fold change of transcripts from Treg treated CD4 (left panel) and CD8 (right panel) Tcon compared to untreated Tcon. Vertical dashed lines on volcano plots indicate a Log2 fold change of 1.5; horizontal dashed line indicates an adjusted p-value of 0.05.

### Treg underwent clonal restriction and activation during GvHD suppression

We next performed the same integrated analysis of the TCR repertoire and the transcriptome on Treg during GvHD suppression. Similar to what we observed in Tcon, Treg underwent clonal restriction during GvHD suppression, as revealed by a significantly increased TCRα and TCRβ clonality index at day 8 compared to before injection (Figure 3A). Accordingly, we observed only limited TCR overlap between day 0 and day 8 Treg (Figure 3B-C). These results indicated that during GvHD suppression Treg underwent clonal restriction at a similar extent as CD4 Tcon (figure 1B). To assess whether Treg and CD4 Tcon reacted to the same antigens during GvHD, we next compared the TCR repertoire of these two subpopulations. We observed a small clonal overlap between Treg and CD4 Tcon before injection and this was further reduced at day 8 after transplantation (Figure 3D-E), suggesting that Treg and CD4 Tcon responses during GvHD are engaging different cell clonotypes triggered by different epitopes or antigens. The increased activation state of Treg during GvHD suppression was further supported by the transcriptomic analysis revealing down-regulation of genes characterizing naive Treg (*Sell*) and up-regulation of several genes involved in activation such as *Icos, Tnfrsf4* (encoding the costimulatory molecule OX40), *Ccr2*, *Klrg1* and *Gzmb* (Figure 3F). After transplantation, Treg preserved the distinct transcriptomic signature observed before injection (Supplemental Figure 3A) further enhanced by the up regulation of genes involved in Treg activation and suppressive function (*Ccr4*, *Ccr8*, *Gata3*, *Il9r*, *Il2ra*, *Il10*, *Tnfrsf18*, *Tnfrsf4*, *Areg*) (Supplemental Figure 3B). Collectively, these data indicate that during GvHD suppression Treg undergo activation and clonal restriction similarly to what was observed in CD4 Tcon, although the analysis of the TCR repertoire of the two populations indicated divergence rather than increased similarity between Treg and CD4 Tcon during GvHD.

**Figure 3.**
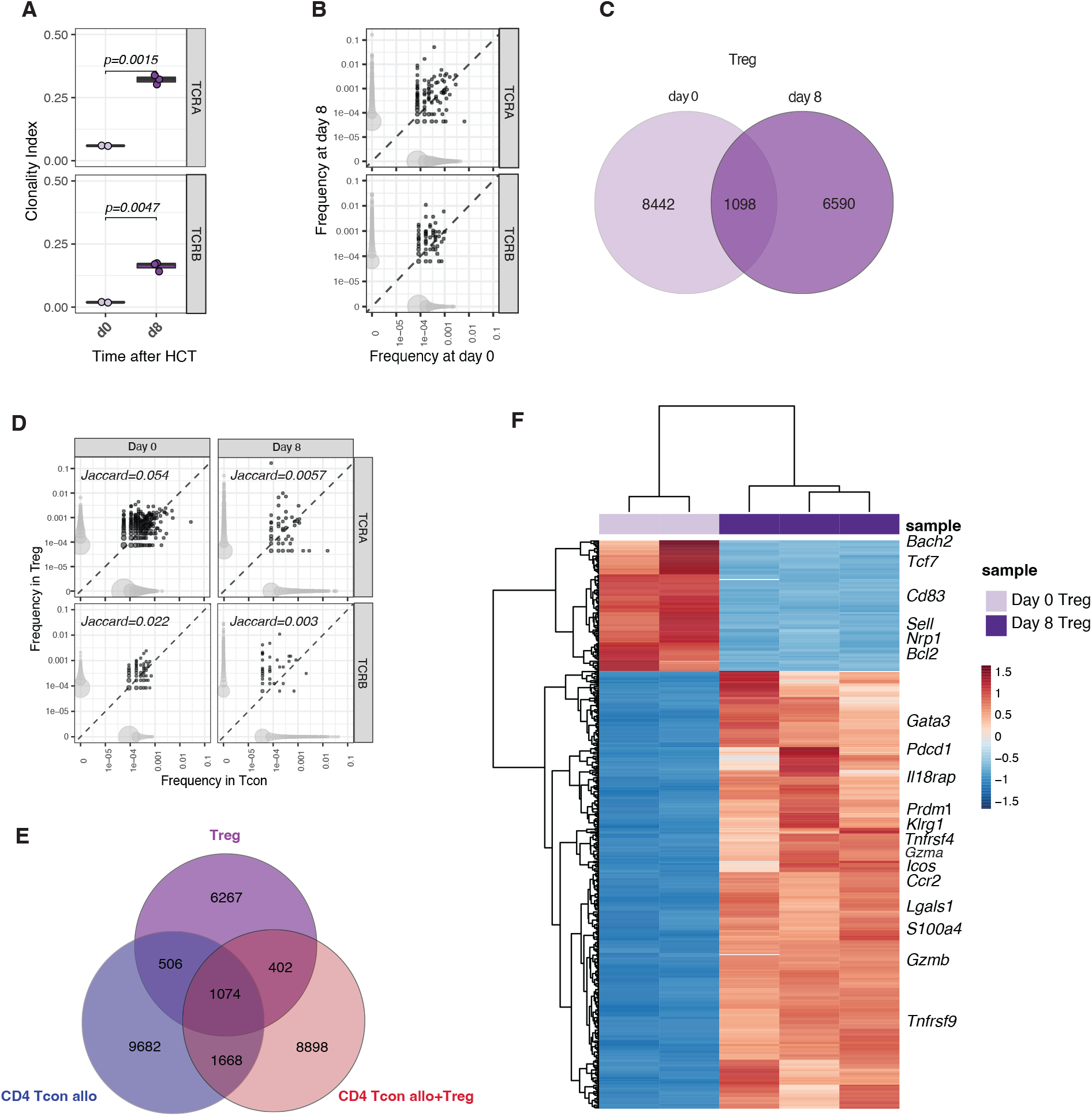
Treg underwent clonal restriction of the TCR repertoire and activation during GvHD suppression. (A) Clonality of the TCRA and TCRB repertoire in Treg at day 0 (light pink box and symbols) and day 8 (purple box and symbols) after HCT. (B) Representative example of overlap of the TCRA and TCRB repertoire in Treg prior to transplantation and at day 8 after HCT. Scatter plots represent clones’ frequencies before and after HCT and number of unique clones (dot size). Clones that are observed at only one time point are colored in light grey, while overlapping clones are colored in dark grey. (C) Venn diagram representing the number of overlapping and non-overlapping clones between day 0 (light pink box and symbols) and day 8 (purple box and symbols) Treg. (D) Representative example of overlap of the TCRA and TCRB repertoire in CD4 Tcon (x axis) and Treg (y axis) at day 8 after HCT in allogeneic mice receiving both Tcon and Treg. Scatter plots represent clones’ frequencies in Tcon and Treg and number of unique clones (dot size). Clones that are observed in only one population are colored in light grey, while overlapping clones are colored in dark grey. Jaccard indexes are indicated. (E) Venn diagram representing the number of overlapping and non-overlapping clones between Treg and Tcon treated or not with Treg recovered at day 8 after HCT. (F) Heatmap and hierarchical clustering based on the 500 most highly differentially expressed genes across all samples. Immune-related genes are highlighted. Expression for each gene is scaled (z scored) across single rows.

### Paired transcriptomic analysis of Treg and Tcon identified IL-10 and IL-35 as potential mechanisms of GvHD suppression

Treg employ a wide range of mechanisms to suppress immunopathological processes, ranging from production of immunosuppressive molecules to metabolic modulation of target cells^21,22^. To infer the dominant mechanisms of suppression employed by Treg to control GvHD from transcriptomic data, we analyzed the transcript expression of suppressive molecules in Treg before and after HCT as well as the expression of gene sets induced by such molecules in Tcon. We did not observe any differences in *Tgfb* gene expression between Treg before injection (day 0) and Treg recovered at day 8 after HCT (Figure 4A). Accordingly, Gene Set Enrichment Analysis (GSEA) of TGFβ-induced genes did not reveal any differences between Tcon treated or not with Treg (Figure 4B). Conversely, Treg at day 8 after HCT expressed higher transcript levels of *Il10* and *Ebi3*, encoding for one of the two subunits constituting IL-35, compared to Treg before injection (Figure 4A). Accordingly, GSEA revealed a significant enrichment of IL-10-induced genes in CD4 Tcon and IL-35-induced genes in CD4 and CD8 Tcon (Figure 4B). Finally, day 8 Treg displayed a significant upregulation of the *Il2ra* gene, encoding the alpha chain of the Il-2 receptor (Figure 4A). In addition to being a constitutively expressed marker in Treg that is further upregulated during activation, IL-2RA expression is also essential in IL-2 deprivation of Tcon, an additional Treg mechanism of suppression^23,24^. GSEA for IL-2 induced genes in Tcon did not reveal any significant difference between Tcon recovered from mice treated or not with Treg (Figure 4B). Collectively, our transcriptomic results support the involvement of IL-10 and IL-35 production by Treg and their downstream signaling in Tcon as major mechanisms of suppression of GvHD by Treg.

**Figure 4.**
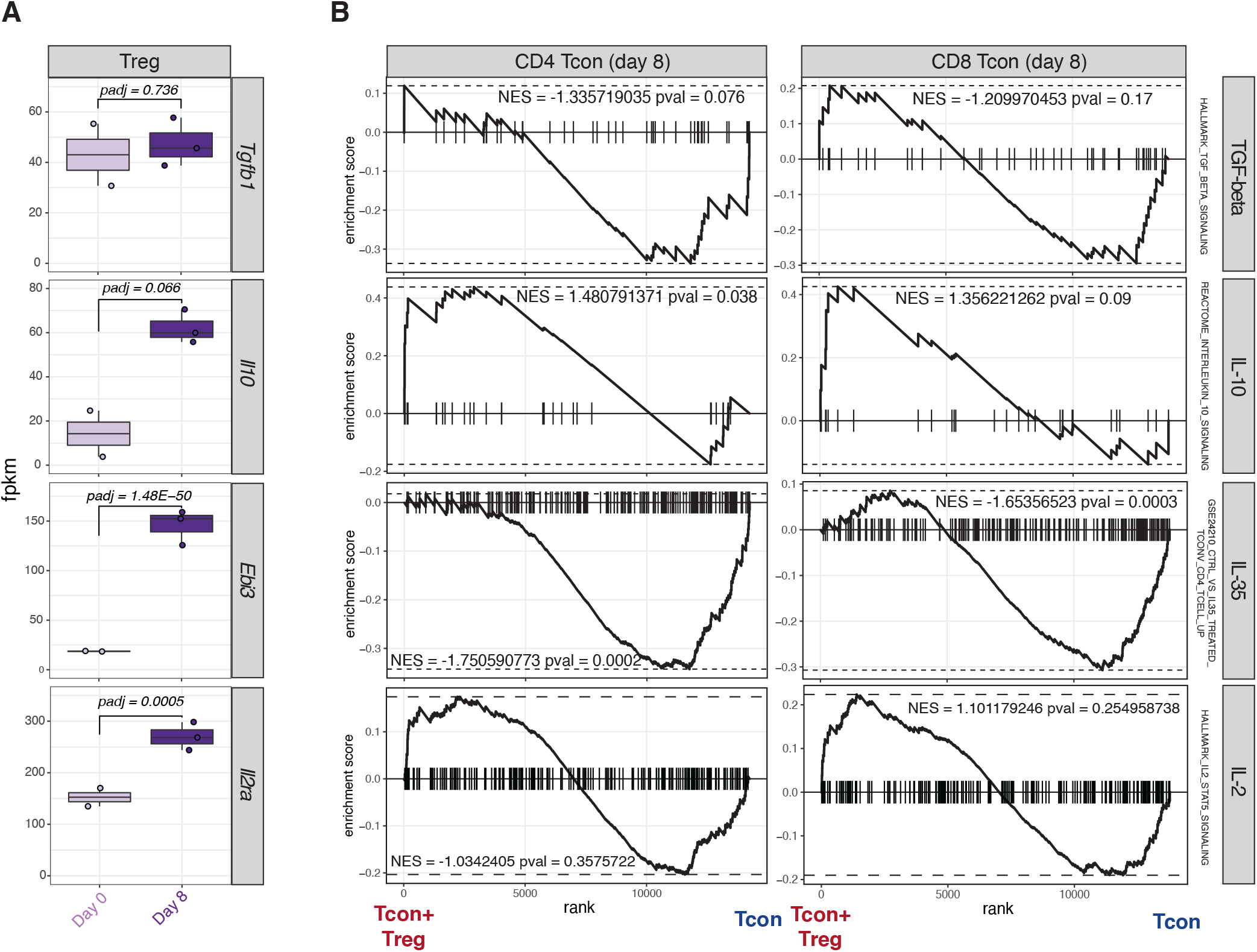
Paired transcriptomic analysis of Tcon and Treg identified mechanisms of GvHD suppression. (A) Transcript expression of genes encoding for molecules involved in classical Treg mechanisms of suppression. (B) Enrichment plots displaying enrichment scores for the genes involved in *Tgfb* (HALLMARK_TGF_BETA_SIGNALING), *Il10* (REACTOME_INTERLEUKIN_10_SIGNALING), *Il35* (GSE24210_CTRL_VS_IL35_TREATED_TCONV_CD4_TCELL_UP) and *Il2* (HALLMARK_IL2_STAT5_SIGNALING) in CD4 (left panels) and CD8 (right panels) Tcon recovered at day 8 after HCT in the presence or absence of Treg. Gene signatures were obtained from Molecular Signatures Database (MSigDB).

### Treg modulated genes involved in metabolic pathways favoring oxidative phosphorylation and suppressing glycolysis in CD4 and CD8 Tcon

To look for additional mechanisms of Treg suppression during GvHD, we performed GSEA for Hallmark gene sets on CD4 and CD8 Tcon in the presence or absence of Treg. This analysis identified the oxidative phosphorylation gene set as the top one induced in CD4 Tcon (Figure 5A) and the top third in CD8 Tcon (Figure 5B). Given the recently discovered importance of T cell-metabolism in GvHD^25–27^, we analyzed in detail the impact of Treg on genes involved in the main metabolic pathways (Figure 5C). Treg treatment significantly suppressed the transcription of genes involved in glycolytic processes including *Slc2a1*, encoding for the glucose receptor GLUT1, and *Pkm*, encoding for the key glycolytic enzyme pyruvate kinase, in both CD4 and CD8 Tcon (Figure 5C). Conversely, Treg treatment led to a global up-regulation of genes encoding for enzymes involved in oxidative phosphorylation (OXPHOS; Figure 5C). We did not observe any significant impact of Treg on Tcon transcription of genes involved in fatty acid oxidation (FAO) or the tricarboxylic acid (TCA) cycle (Figure 5C). Collectively, our results demonstrated that Treg significantly modulated genes involved in Tcon metabolism, leading to the downregulation of genes involved in glycolysis and the upregulation of the ones responsible for oxidative phosphorylation, pointing to a metabolic shift of Tcon induced by Treg during GvHD.

**Figure 5.**
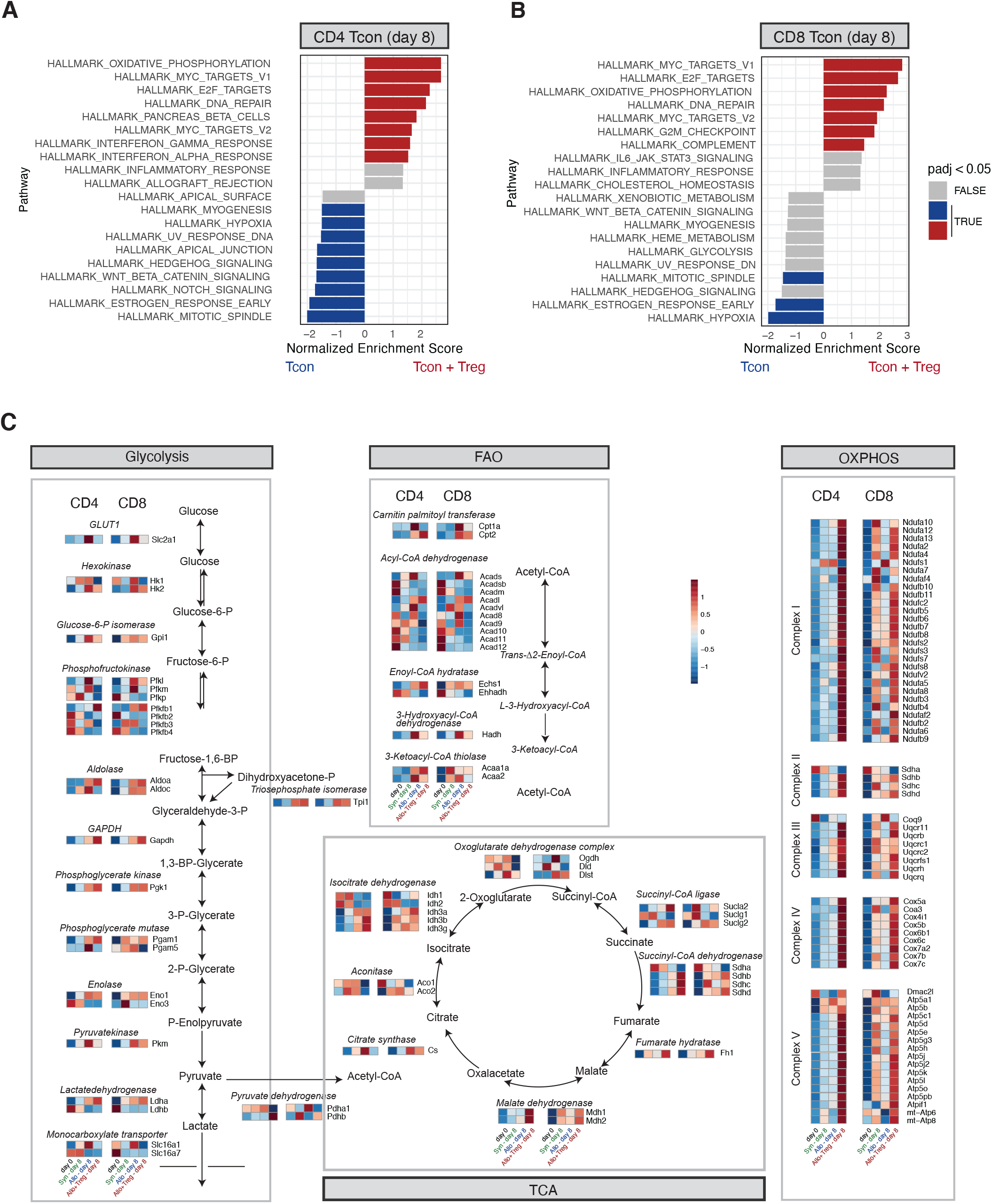
Treg modulated genes regulating metabolic patterns in CD4 and CD8 Tcon during GvHD. (A-B) Top 10 enriched terms/pathways in CD4 (A) and CD8 (B) Tcon from mice receiving Tcon and Treg (positive NES, red bars indicate the significant patways) or Tcon alone (negative NES, blue bars indicate the significant pathways) revealed by Hallmark GSEA. (C) Map of genes regulating CD4 and CD8 Tcon metabolisms before and after HCT. Single genes heatmap represent the row-scaled gene expression expressed in FPKM. Genes and enzymes are indicated.

### Treg treatment did not affect the induction of effector gene sets involved in GvT effect

We and others have previously shown that Treg are capable of suppressing GvHD without impairing the GvT effect of the transplant procedure^5^. We therefore hypothesized that Treg treatment would have minimal if any impact on Tcon transcription of effector molecules involved in the GvT effect. To test this hypothesis, we compared the transcription of effector molecules in Tcon recovered at day 8 from allogeneic mice treated or not with Treg. As shown in Figure 6A, the addition of Treg did not inhibit but further increased the transcription of *Ifng*, *Il2* and *Tnf* in CD4 Tcon. Accordingly, Treg did not prevent the upregulation of gene sets involved in leukocyte mediated cytotoxicity after HCT (Figure 6B). Similar results were observed in CD8 Tcon where Treg did not prevent the upregulation of cytotoxic genes including *Ifng*, *Gzmb* and *Prf1* (Figure 6C), nor the enrichment in genes involved in leukocyte mediated cytotoxicity (Figure 6D). Collectively, these results demonstrate that Treg treatment did not interfere with the induction of genes encoding effector molecules involved in the GvT effect of HCT.

**Figure 6.**
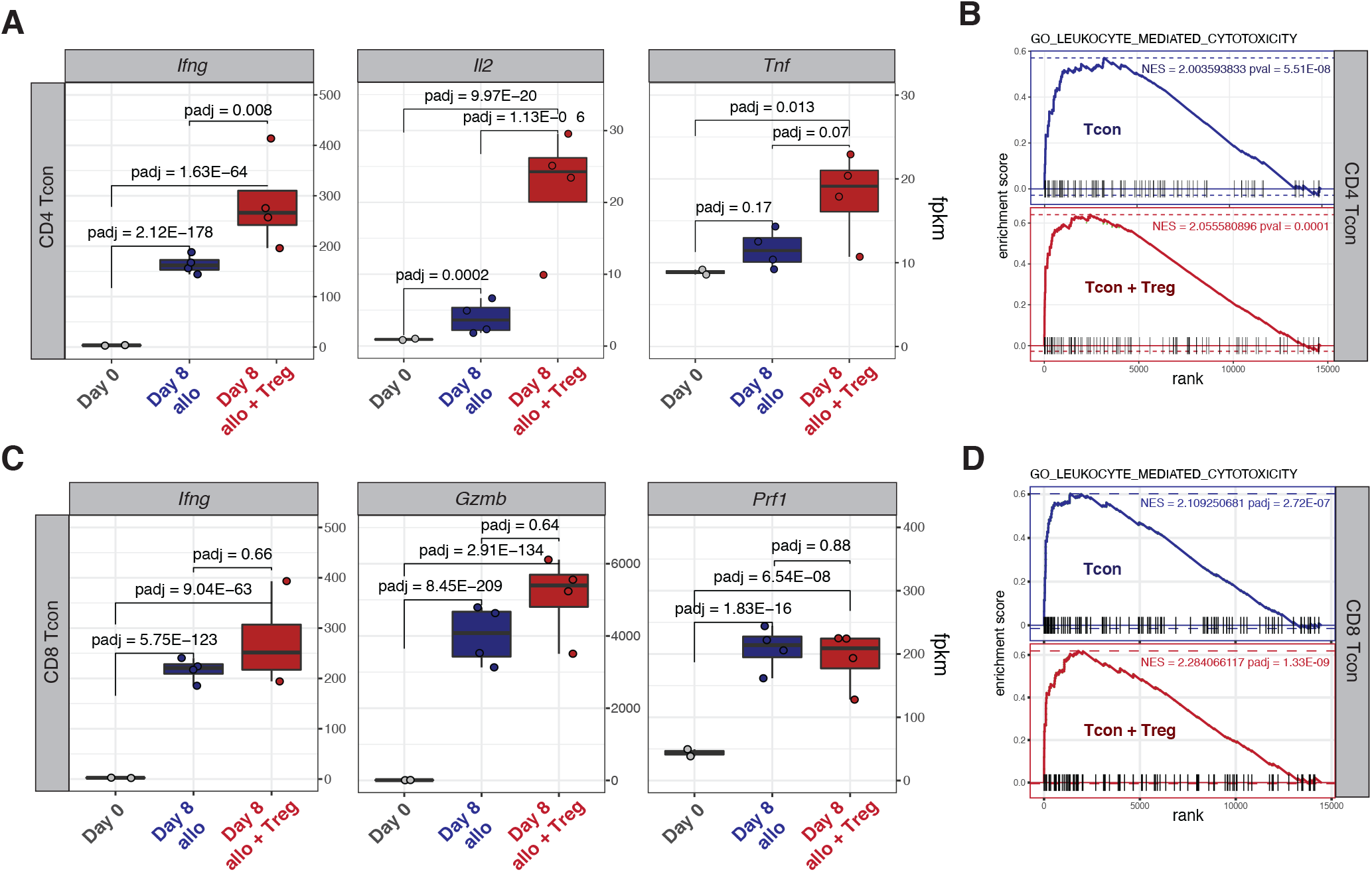
Treg did not inhibit the upregulation of gene sets involved in the GvT effect. (A) Transcript expression of effector molecules (*Ifng*, *Il2*, *Tnf*) in CD4 Tcon. (B) Enrichment plots displaying enrichment scores for the genes involved in leucocyte cytotoxicity (GO_LEUKOCYTE_MEDIATED_CYTOTOXICITY) from CD4 Tcon recovered at day 8 after HCT in the presence or absence of Treg. (C) Transcript expression of effector molecules (*Ifng*, *Gzmb*, *Prf1*) in CD8 Tcon. (D) Enrichment plots displaying enrichment scores for the genes involved in leucocyte cytotoxicity (GO_LEUKOCYTE_MEDIATED_CYTOTOXICITY) from CD8 Tcon recovered at day 8 after HCT in the presence or absence of Treg. Gene signatures were obtained from Molecular Signatures Database (MSigDB).

## Discussion

In this work we used integrated TCR repertoire and transcriptomic analysis of murine Tcon and Treg to gain further insights into the mechanisms of acute GvHD suppression by Treg. Our results indicate that Treg treatment did not interfere with the activation and differentiation of alloreactive Tcon clones during GvHD. Treg predominantly affected the CD4 Tcon and to a lesser extent the CD8 Tcon transcriptome, modulating the transcription of genes encoding pro- and anti-inflammatory molecules as well as enzymes involved in glycolytic processes.

CD4+CD25+FOXP3+ regulatory T cells are a well-established immunomodulatory cell population able to suppress conventional T cell responses employing several, non-mutually exclusive, mechanisms^21^ including: 1) production of immunosuppressive molecules, namely TGFβ, IL-10 and IL-35; 2) metabolic disruption by IL-2 competition and pericellular generation of adenosine; 3) cytolytic activity toward activated T cells or antigen presenting cells (APC); 4) modulation of APCs. Our analysis identified multiple pathways potentially involved in Treg suppression of GvHD, namely anti-inflammatory cytokine production mirrored by downstream Tcon signaling of IL-10 and IL-35 as well as a metabolic switch of Tcon from glycolysis to oxidative phosphorylation. Conversely, we did not find evidence for a role of TGFβ production and competition for IL-2 as dominant mechanisms of Treg suppression. The study of Tcon and Treg transcriptome during GvHD limited our analysis to T cell intrinsic mechanisms of suppression while it did not allow us to evaluate the relevance of cytolysis of effectors and APCs. Previous studies addressing the role of cytolysis mediated by Treg through production of cytotoxic molecules failed to find evidence for a role of granzyme B^28^ and showed experimental^29^ and clinical^30^ evidence for a role of granzyme A in Treg mediated suppression of GvHD. Our transcriptomic analysis found an upregulation of *Gzmb* and *Gzma* in Treg after transplantation (Figure 3F).

Recent studies point to an important role of metabolic regulation of T cells during GvHD (reviewed in *Mohamed et al*^31^). Murine studies revealed that donor T cells undergo metabolic reprogramming after allogeneic HCT, switching from fatty acid β-oxidation (FAO) and pyruvate oxidation via the tricarboxylic (TCA) cycle to aerobic glycolysis^25^. Using transcriptomic and metabolomic analysis, Assmann and colleagues^26^ confirmed that murine donor CD4+ T cells acquired a highly glycolytic profile during acute GvHD and showed increased transcription of glycolytic enzymes in human CD4+ T cells isolated from allogeneic HSCT recipients just before the onset of acute GvHD. Our transcriptomic results suggest that Treg inhibit the metabolic switch of Tcon toward glycolysis by interfering at different crucial points. We observed a decrease in the transcription of the gene encoding the glucose transporter GLUT1, which contributes to the pathogenicity of allogeneic Tcon during GvHD^27,32^. Moreover, Treg inhibited the induction of genes encoding several glycolytic enzymes, including 6-phosphofructo-2-kinase/fructose-2,6-biphosphatase 3 (PFKFB3), the rate-limiting factor in glycolytic metabolism whose specific pharmacological inhibition using 3-(3-pyridinyl)-1-(4-pyridinyl)-2-propen-1-one (3-PO) has been shown to protect against acute GvHD^25^. T cell metabolic fitness through glycolysis and oxidative phosphorylation (OXPHOS) has been recently shown to play an essential role in the GvT effect after allogeneic HCT^33^. In our experiments, Treg not only inhibited the transcription of glycolytic genes but also increased the transcription of OXPHOS-related genes, suggesting a metabolic switch toward mitochondrial respiration as a source of energy.

The need for TCR activation as well as the nature of the antigens recognized by Treg during GvHD is still debated. The beneficial effects of low dose IL-2 treatment on Treg numbers and function in chronic GvHD^34–36^ suggest that cytokine-mediated Treg activation is sufficient for GvHD suppression without the need for TCR triggering. However, we previously showed that MHC disparities between Treg and the host were necessary as both donor and third-party Treg but not host Treg protect from GvHD in murine allogeneic HCT^37^ pointing to a critical role of TCR activation for alloreactive Treg suppression of GvHD. Our present study further supports a model of TCR-mediated Treg activation for GvHD suppression, given the clonal restriction that Treg undergo after allogeneic HCT. Interestingly, we observed a divergence rather than a convergence of Tcon and Treg clonotypes detected after HCT compared to the steady state, suggesting that Tcon and Treg react against different antigens during GvHD. The contribution of Tcon derived IL-2 to this process is not excluded. Surprisingly, IL-2 production by CD4 Tcon was increased rather than inhibited by the presence of Treg, potentially providing an additional source of cytokines to support Treg homeostasis and function.

Our results have clinical implications given the increasing interest in Treg-based therapies for GvHD prevention and treatment. The Perugia group pioneered the adoptive transfer of fresh Treg followed by Tcon in a T cell depleted, CD34-selected HLA-haploidentical HCT platform^7,10,38^ demonstrating the potential of human Treg to prevent GvHD but to still allow the GvT effect in patients. We reported a similar approach in HLA-matched recipients^9^. Recently, therapeutic adoptive transfer of Treg-enriched donor lymphocyte infusion combined with low-dose IL-2 has been reported in chronic GvHD^11^. A better understanding of the mechanisms of GvHD suppression by Treg is particularly relevant now that, after years of monocentric early-phase clinical trials, the field is moving to the first multicentric phase III clinical trials.

Given the rarity of Treg, several groups attempted to *ex vivo* expand them from cord-blood^8,40^ or from peripheral blood^12^. Our results point to the need of optimizing culture conditions to favor the expansion of IL-10 and IL-35 producing Treg^41^.

Our data reveal that, during GvHD suppression, Treg preserved a transcriptomic signature distinct from CD4 and CD8 Tcon (Supplemental Figure 3). Among differentially expressed genes, we identified genes encoding several surface markers, including Killer cell lectin-like receptor family molecules (*Klrc1*, *Klrd1*, *Klrk1*, *Klrb1b*), *CD160* and Cytotoxic and regulatory T cell molecule (*Crtam*). Efforts are ongoing to target these and other markers to selectively deplete alloreactive T cells while sparing Treg.

Our study has several limitations. First, we performed our analysis on Treg and Tcon recovered from secondary lymphoid organs (SLO) and not on GvHD-target tissues as previously reported by other groups who focused on Tcon^42–45^. We decided to study SLO because this is the site where Treg suppression of Tcon mediating GvHD is believed to take place^6,46^. Moreover, the reduction in Tcon tissue infiltration upon Treg treatment precluded this kind of analysis at the GvHD-target tissues sites. Second, our analysis was performed at the peak of Tcon expansion and lacks the dynamic information about the early impact of Treg during the very first days of GvHD suppression. Unfortunately, the limited number of cells that is possible to recover at earlier time points represent an obstacle to this analysis. Third, the strain combination of C57Bl/6 donors into BALB/c recipients is known to be more dependent on CD4+ T cells where it is possible that other strain combinations more dependent on CD8+ T cells may show more impact of Treg on this cell population.

In conclusion, our results provide further insights into the mechanisms of Treg suppression of GvHD. Moreover, our data support a model in which Treg qualitatively modulate Tcon function through several mechanisms rather than preventing the activation of alloreactive clones, providing a potential explanation for the ability of Treg to suppress GvHD while allowing GvT.

## Author contributions

JKL, TH, MT, FS conceived and designed research studies; JKL, TH, MT, TLR, PYL, FS conducted experiments; JKL, SB, NK, FS analyzed data; XJ, developed methodology and analyzed data; JB, developed methodology and provided essential reagents; JV, YC provided essential tools and intellectual input; JKL, FS and RSN wrote the manuscript; FS and RSN supervised the research

## Data and materials availability

Relevant data can be found in the Gene Expression Omnibus database (accession number GSE205375; https://www.ncbi.nlm.nih.gov/geo/query/acc.cgi?acc=GSE205375).

## Supplemental figure legends

**Supplemental Figure 1.**
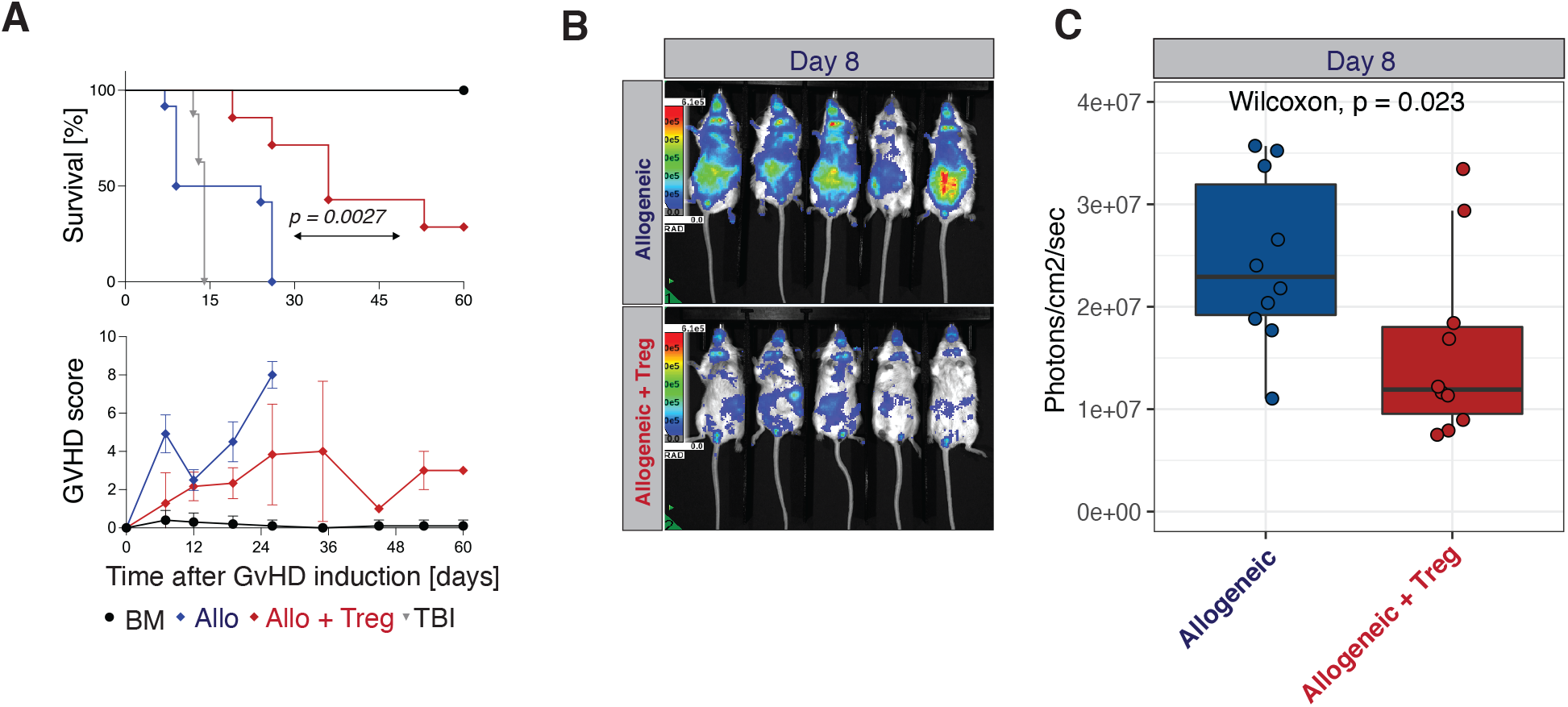
Treg extended survival, reduced clinical signs and inhibited Tcon expansion and localization to target organs in a murine model of acute GvHD. (A) Survival (upper panel) and GvHD score (lower panel) of lethally irradiated mice (grey), transplanted with BM cells alone (black), BM plus allogeneic Tcon (blue) or BM plus allogeneic Tcon and Treg (red). (B) Representative bioluminescence (BLI) images (B) and analysis (C) of mice receiving allogeneic Tcon (upper panel) or allogeneic luciferase^+^ Tcon together with Treg (lower panel). Results are pooled from two independent experiments with a total of 10 mice per group. Values were compared using a nonparametric Mann-Whitney test. P value is indicated.

**Supplemental Figure 2.**
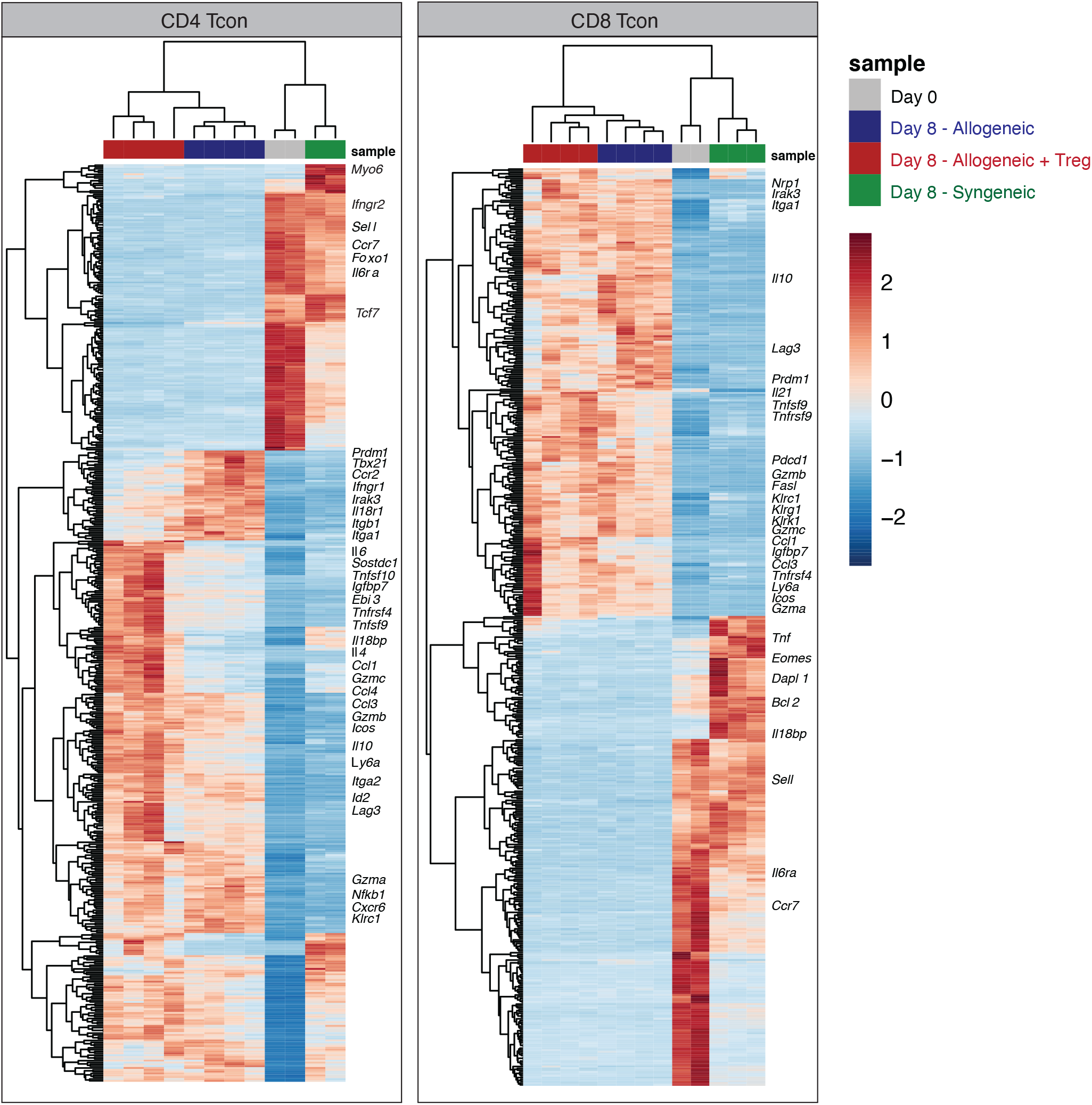
Treg modulated the transcription of multiple immunological relevant genes in CD4 and CD8 Tcon at day 8 after HCT. Heatmap and hierarchical clustering based on the 500 most highly differentially expressed genes across all samples. Immune-related genes are highlighted. Expression for each gene is scaled (z scored) across single rows.

**Supplemental Figure 3.**
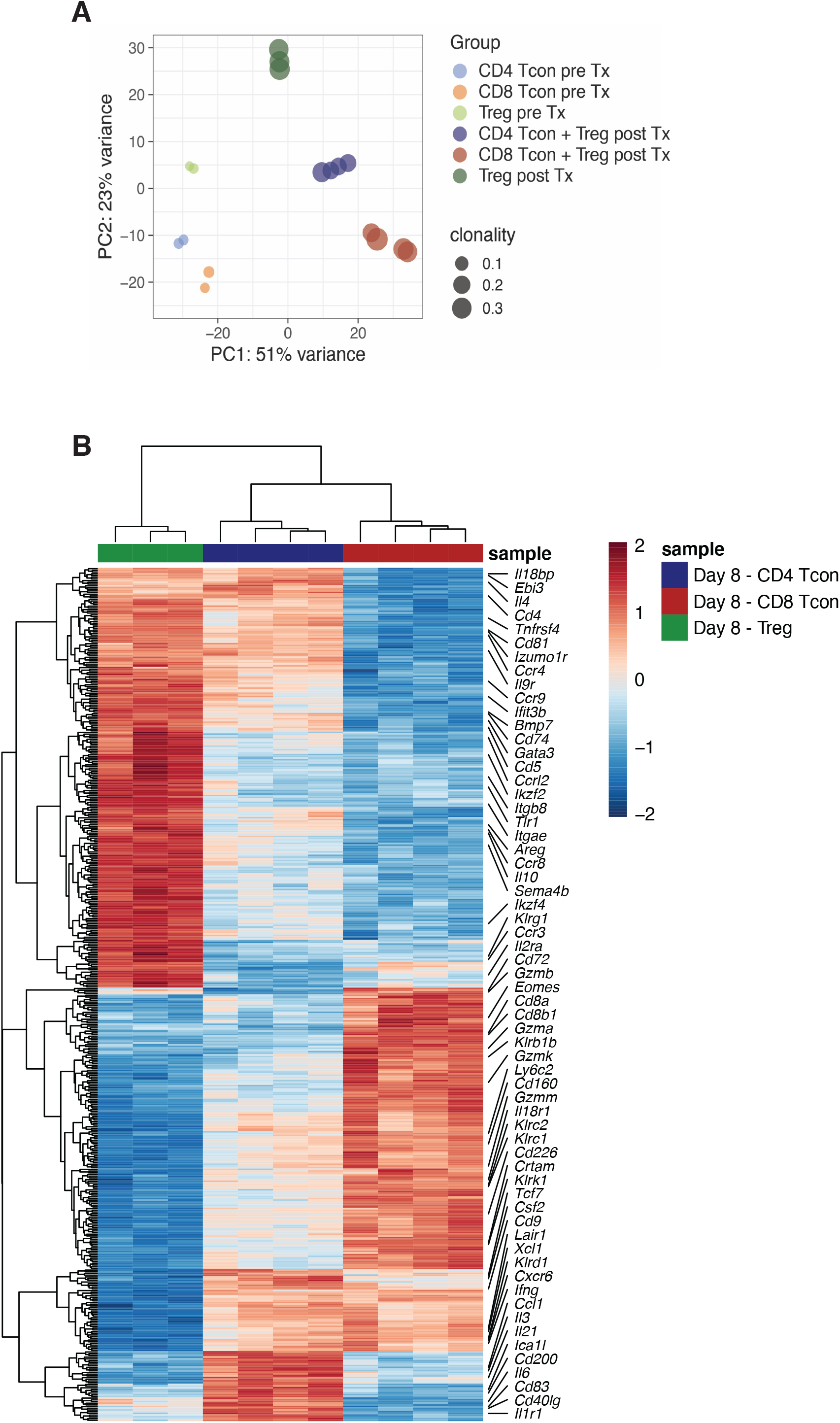
Treg preserved a transcriptomic signature distinct from CD4 and CD8 Tcon at day 8 after HCT. (A) Principal component analysis (PCA) and clonality of Treg, CD4 and CD8 Tcon before injection and recovered at day 8 post-HCT from Treg treated mice. PCA is based on the 1000 most differentially expressed genes. TCR alpha clonality index is represented. Pre Tx: before transplantation (day 0); post Tx: after transplantation (day 8). (B) Heatmap and hierarchical clustering based on the 500 most highly differentially expressed genes across all samples. Immune-related genes are highlighted. Expression for each gene is scaled (z scored) across single rows.

